# BioFuse: An embedding fusion framework for biomedical foundation models

**DOI:** 10.1101/2025.03.01.640976

**Authors:** Mirza Nasir Hossain, David Harris-Birtill

## Abstract

The biomedical field has witnessed a surge in pre-trained foundation models excelling in specific sub-domains such as radiology and histopathology. While integrating these models promises a more comprehensive understanding of biomedical data, it poses challenges in model compatibility and feature fusion. We present BioFuse, a novel open-source framework designed to generate optimised biomedical embeddings. BioFuse utilises a pool of 9 state-of-the-art foundation models to create task-specific embeddings. It employs grid search to automatically identify the optimal combination of models, fusing their embeddings through vector concatenation. On the MedMNIST+ benchmark, using XGBoost as the downstream classifier, BioFuse outperforms several existing methods, achieving SOTA AUC in 5/12 datasets while maintaining near-SOTA performance across most remaining datasets. Remarkably, our experiments reveal unexpected cross-modal capabilities, with histopathology and radiology models showing strong performance when applied to other imaging modalities. BioFuse features a high-level API^1^ for immediate deployment and an extensible architecture to incorporate future models and fusion techniques. We anticipate BioFuse will not only enhance the utility of foundation models in biomedicine but also open new avenues for uncovering cross-modal relationships in biomedical data.

## Introduction

Artificial Intelligence (AI) is driving advances in biomedical research and clinical practice [1–3]. Foundation models — large, pre-trained models capable of performing a wide variety of tasks with minimal adaptation — have emerged as key tools in this domain. Trained on vast, diverse datasets [4], these models have demonstrated promising performance across various biomedical sub-domains [5].

In medical imaging, foundation models are utilised for detecting and classifying abnormalities in radiographs [5], generating diagnostic reports [6], segmenting images to delineate organs and tumours [7], analysing histopathology slides for disease diagnosis [8, 9], and integrating information across imaging modalities to enhance diagnostic accuracy [10]. In clinical natural language processing (NLP), they are employed for clinical text summarization [11] and information extraction [12], while in genomics, they assist in deciphering the language of non-coding DNA [13] and enable accurate cell type annotation [14].

Typically, researchers manually select foundation models tailored for specific tasks [15]. These models increasingly serve as feature extractors, transforming input data into dense numerical representations called embeddings [16]. These embeddings encapsulate complex features of the input data in high-dimensional vectors, allowing for efficient computation and comparison.

However, foundation models are often trained on single modalities, with research groups developing specialized architectures for specific domains [17–19]. While this approach enables exceptional performance within designed scopes, these models may have learned features relevant beyond their original domains. As domain-specific models become increasingly refined [20], there’s a growing need to explore their broader applicability across biomedical tasks and modalities.

Combining embeddings from multiple models allows us to exploit modality-specific encodings while gaining new perspectives through cross-modal integration. Various approaches have demonstrated the value of leveraging multiple modalities for instance, ConVIRT [21] utilises contrastive learning between paired images and text to enhance chest X-ray classification performance. Such cross-modal techniques suggest that strategic integration of specialized models could generate more comprehensive representations of biomedical data and unlock insights inaccessible to single-modality approaches.

This leads us to ask two key questions:

1. Does combining the embeddings of multiple foundation models trained on different biomedical modalities improve task performance, as measured by AUC (area under the curve) and accuracy?
2. How effectively do foundation models transfer knowledge across different biomedical imaging modalities, as evidenced by comparative AUC and accuracy scores in cross-modal applications?

Despite the potential benefits, effectively integrating multiple foundation models presents major challenges. Retraining models on combined data from multiple modalities demands extensive computational resources and expertise [22]. While combining pre-trained model embeddings offers an alternative [23, 24], the rapidly growing number of foundation models [25] makes identifying optimal combinations increasingly complex. Moreover, the lack of standardized frameworks for embedding fusion hinders our ability to fully exploit these models’ collective knowledge.

### Our Contributions

In response to these challenges, we introduce BioFuse, an open-source framework designed to harness the collective power of diverse biomedical foundation models. This paper makes the following key contributions:

1. **Enhanced Performance on the MedMNIST+ Benchmark:** Using embeddings generated by BioFuse combined with XGBoost classification, we outperform several existing baselines on the MedMNIST+ benchmark, achieving the highest test AUC in 5 of 12 datasets and best accuracy in 2 datasets. The framework’s embeddings maintain near-SOTA performance (within 2% margin) across most remaining datasets, demonstrating the effectiveness of our embedding fusion approach.
2. **Revealing Cross-Modal Capabilities in Biomedical Foundation Models:** Our experiments reveal unexpected cross-modal transfer in both histopathology and radiology models, showcasing the potential for models trained on one imaging modality to generate effective embeddings for others.
3. **An Extensible Framework for Optimised Biomedical Embedding Generation:** BioFuse provides a high-level API that simplifies the generation of task-specific embeddings. It automatically identifies optimal combinations of foundation models and fuses their outputs, addressing the challenges of model integration. The framework’s architecture allows for easy incorporation of new models and fusion techniques, ensuring adaptability to emerging developments in biomedical AI.

By providing a standardized approach to embedding fusion and automating model selection, BioFuse enhances the utility of foundation models in biomedicine while opening new avenues for uncovering relationships between diverse biomedical data representations.

## Background

### Foundation Models in Biomedicine

Foundation models, driven by Large Language Models (LLMs), have ushered in a new era in deep learning [26]. Three key factors enabled their development: the Transformer architecture [27] for processing sequential data, advances in GPU capabilities [28], and extensive training datasets [29, 30]. These billion-parameter models [12] require substantial computing infrastructure for training [31], but can then solve tasks beyond their original training objectives [32]. While they support zero-shot inference and finetuning [26], they are commonly used as feature extractors for classification models [16, 33, 34].

Foundation models are typically trained using self-supervised learning (SSL) techniques, including Contrastive Learning [35], Masked Image Modelling (MIM) [36], and self-distillation [37]. These methods create proxy tasks from unlabelled data: contrastive learning differentiates between similar and dissimilar samples, MIM reconstructs masked image regions, and self-distillation leverages the model’s own predictions as training targets. By utilizing the inherent structure of data, SSL enables training on much larger datasets without manual annotation. This approach has demonstrated superior performance and transferability compared to traditional supervised learning [35, 36, 38], making it particularly valuable in medical imaging where labelled data is scarce.

### Unimodal Foundation Models

Biomedical imaging foundation models have demonstrated remarkable effectiveness when trained on single modalities (unimodal training). In computational pathology, models like UNI [39], UNI2 [40], Prov-GigaPath [41], and Hibou-B [42] have set new benchmarks through increasingly larger-scale training. UNI outperforms prior models across 34 clinical tasks, while its successor UNI2 expands capabilities to both H&E and Immunohistochemistry (IHC). Prov-GigaPath achieves state-of-the-art performance in cancer subtyping and mutation prediction, and Hibou-B demonstrates high adaptability across pathology applications. In radiology, RAD-DINO [43], pre-trained on a vast corpus of chest X-rays, excels in detecting conditions and capturing detailed features crucial for biomarker discovery and prognosis prediction. Built on architectures like Vision Transformers (ViT) and trained with self-supervised learning, these models demonstrate how large-scale training captures modality-specific nuances and enables superior biomedical imaging performance.

### Vision Language Models

Vision-language models (VLMs) integrate visual and textual data to learn joint representations for tasks like image classification, retrieval, and captioning. BioMedCLIP [44] and PubMedCLIP [45] achieve state-of-the-art performance across various biomedical benchmarks, while CheXagent [46] excels in chest X-ray interpretation tasks and CONCH [47] demonstrates superior performance across 13 histopathology benchmarks. These models demonstrate how integrating visual and textual information enhances biomedical imaging analysis.

### Limitations of Foundation Models

Despite their capabilities, biomedical foundation models face serious limitations including restricted cross-domain applicability, high computational demands, and challenges in interpretability crucial for clinical decisionmaking. While using them as feature extractors mitigates the risk of hallucinations [20], effectively leveraging the collective knowledge of multiple specialized models remains an open challenge that could enhance performance and provide more robust biomedical insights.

### Related Work

Prior work relevant to BioFuse spans three key areas: embedding fusion approaches that combine features from multiple models, cross-modal methods that transfer knowledge between domains, and automated selection techniques that optimize model combinations.

### Embedding Fusion

Recent works have demonstrated the benefits of combining embeddings from multiple models. In histopathology, Neidlinger et al. [24] showed that combining four foundation models improved tumor classification and survival prediction across 13 patient cohorts, but their approach was limited to a single modality and lacked a framework for optimal model selection.

Similarly, Zarif et al. [48] and Dong et al. [49] combined two and four pre-trained CNNs respectively for breast and liver cancer classification, but their approaches used bespoke architectures specific to their tasks. In clinical NLP, BioFLAIR [50] combines two embedding models (FLAIR and BioELMo) for biomedical named entity recognition, while the Concatenated BioMed-Transformers [51] fuses three transformer models for medical text classification, but both were limited to single modalities.

### Cross-Modal Transfer

Cross-modal approaches have attempted to bridge different modalities in biomedical data. Zipper [52] combines two pre-trained unimodal models for speech recognition and text-to-speech tasks using multi-tower decoders with cross-attention, but its computational complexity limits practical application. BioBridge [53] uses knowledge graphs to connect two unimodal foundation models for protein sequence-text retrieval and drug design tasks, yet fails to fully exploit the embedded knowledge across different biomedical domains.

### Automated Model Selection

In automated model selection, ACE [54] uses reinforcement learning to select optimal combinations of word embedding models for NLP tasks like named entity recognition and part-of-speech tagging. However, its focus on text-only tasks fails to address the multimodal nature of biomedical data, and its computational demands make it impractical for high-dimensional data where rapid model selection is often needed.

### Limitations of Related Work

While these approaches have made impressive strides, they often focus on specific domains or tasks, lacking the versatility to leverage image extraction capabilities across models trained on different modalities. Moreover, the untapped potential in combining foundation models to enhance performance and generalisability across diverse modalities remains a non-trivial challenge. BioFuse aims to address these limitations by providing a flexible framework that can integrate multiple foundation models across various biomedical domains, facilitating both embedding fusion and cross-modal transfer while automating the selection of optimal model combinations.

## BioFuse

### Overview

BioFuse is an open-source framework that automates the selection, extraction, and fusion of embeddings from multiple pre-trained biomedical foundation models across diverse modalities such as radiographs, histopathology slides, and clinical text. Its modular design supports seamless integration of new models and fusion methods, keeping pace with rapid advancements. BioFuse’s primary goal is to generate optimal embeddings by leveraging multiple models via vector concatenation. These fused embeddings encapsulate multimodal information in a unified format. To assess their quality, BioFuse employs an approach similar to linear probing [55], but with a more sophisticated classifier. Specifically, we use XGBoost [56], a powerful tree-based machine learning algorithm, to train on the frozen, fused embeddings for various downstream tasks. This evaluation method provides a reliable measure of embedding quality without requiring computationally expensive fine-tuning of the foundation models themselves.

Users can utilise these embeddings as input features for custom models tailored to specific research questions or clinical applications. By offering optimised embeddings and demonstrating their effectiveness through advanced probing, BioFuse serves as a versatile feature extraction framework for biomedical imaging tasks.

### System Architecture

BioFuse’s architecture comprises three components: pre-trained foundation models, the BioFuseModel, and the search module (see Fig 1):

1. **Pre-trained Foundation Models:** Existing, pre-trained models from various biomedical domains are loaded without fine-tuning to preserve their original capabilities.
2. **BioFuseModel:** The core component that processes and integrates embeddings from multiple pre-trained models in a single forward pass per input. It preprocesses each model’s input, extracts embeddings, and fuses them via vector concatenation, preserving the strengths of each model while ensuring efficiency.
3. **Search Module:** Evaluates embedding performance from different model combinations (each represented by a BioFuseModel) by computing validation accuracy using XGBoost as a lightweight downstream classifier. This step automates identifying optimal configurations.

**Fig 1.**
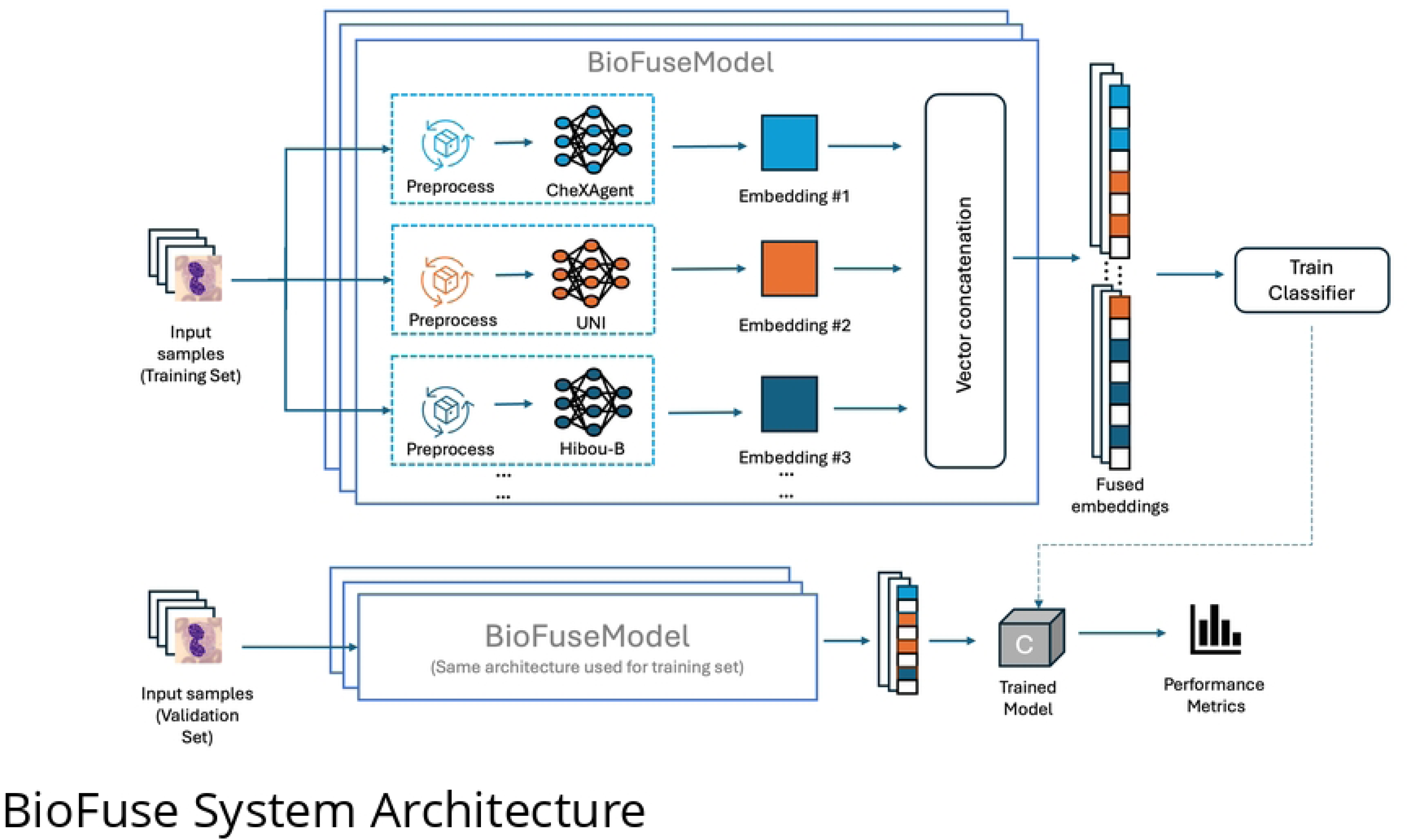
BioFuse architecture and workflow. The upper section illustrates the training process: input samples are preprocessed and fed into multiple foundation models to generate embeddings, which are then concatenated and used to train a classifier. The lower section shows the evaluation process using the same BioFuseModel architecture on a validation set, followed by performance assessment of the trained model.

### Foundation Models in BioFuse

Foundation models in BioFuse were selected based on architectural diversity, performance, efficiency, and accessibility. Both unimodal and vision-language models (VLMs) are included to capture diverse biomedical information, with VLMs chosen for their pre-training on paired text-image data. Models trained on high-quality biomedical datasets in various imaging modalities, from macroscopic radiological images to microscopic histological data, were preferred for a comprehensive representation of knowledge.

The selection process prioritized models documented in peer-reviewed literature with demonstrated excellence across multiple biomedical applications. We carefully balanced computational efficiency against performance to ensure BioFuse remains practical for real-world implementation. Additionally, we favored models with standardized implementations on platforms such as Hugging Face [57] to enhance accessibility and community support. Through this thoughtful curation, BioFuse incorporates a diverse array of foundation models that collectively provide robust and adaptable capabilities across the biomedical domain.

Table 1 summarizes the foundation models supported by BioFuse, including dataset size, pre-training methods, and encoder types.

**Table 1.** Summary of Foundation Models supported in BioFuse.

| Model | Type | Dataset | Dataset size | Pre-training method | Text Encoder | Vision Encoder |
| --- | --- | --- | --- | --- | --- | --- |
| BioMedCLIP [44] | VLM | Figure-caption pairs from PubMed Central | 15M | Contrastive Learning | PubMedBERT | ViT |
| PubMedCLIP [45] | VLM | Radiology Objects in Context (ROCO) | 81K | Contrastive Learning | Transformer | ViT |
| CheXagent [46] | VLM | CheXinstruct (28 CXR datasets) | 6.1M I-Q-A, 1.1M CXRs | Instruction tuning [58], Vision-language pre-training | Custom clinical LLM | ViT |
| CONCH [47] | VLM | Histology slides | 1.17M image-caption pairs | Vision-Language pre-training [32] | CLIP-based | ViT-B/12 |
| rad-dino [43] | Unimodal | MIMIC-CXR, CheXpert, NIH-CHR, PadChest, BRAX | 838K | DINOv2 | - | ViT |
| UNI [39] | Unimodal | Mass-100K | 100M tiles, 100K WSIs | DINOv2 | - | ViT-L/16 |
| UNI2 [40] | Unimodal | 200M H&E and IHC images | 200M tiles, 350K WSIs | DINOv2 | - | ViT-h/14-reg8 |
| Hibou-B [42] | Unimodal | Histology slides | 512M tiles, 1M WSIs | DINOv2 | - | ViT-B/14 |
| Prov-GigaPath [41] | Unimodal | Histology slides | 1.3B tiles, 171K WSIs | DINOv2 | - | ViT |

VLM: Vision-Language Model, ViT: Vision Transformer, I-Q-A: Instruction-Question-Answer

### Embedding Extraction

BioFuse streamlines the extraction of embeddings from diverse pre-trained foundation models across multiple biomedical imaging modalities. The framework leverages established libraries including Hugging Face Transformers [59], OpenCLIP [60], and PyTorch Image Models [61] to facilitate this process. The system automatically handles model and preprocessor loading while applying appropriate model-specific preprocessing—such as image resizing and normalization—before generating embeddings.

The framework employs GPU acceleration to enhance inference speed when processing large-scale datasets. Memory management is optimised through immediate release of intermediate outputs after use, enabling the system to scale effectively with increasingly complex inputs. These technical enhancements ensure BioFuse can efficiently handle diverse biomedical imaging modalities while maintaining computational efficiency in resource-constrained environments.

### Fusion Methodology

BioFuse fuses embeddings from multiple pre-trained models using **vector concatenation**, combining embeddings into a single representation.

Let 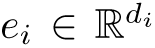 be the embedding from the *i*-th model, where *d_i_* is its dimensionality. Given *n* models, the fused embedding is:

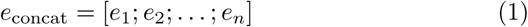

where [;] denotes concatenation along the feature dimension, yielding a total size:

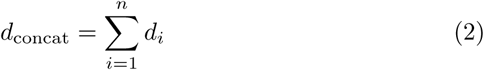

For instance, if 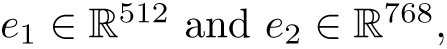 then 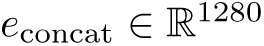.

### Advantages of Vector Concatenation

- **Efficient & Simple**: No additional parameters or projection layers, reducing computational complexity.
- **Feature Preservation**: Retains all modality-specific features without loss.
- **Scalable**: New models can be seamlessly integrated without retraining fusion components.
- **Interpretable**: Maintains per-model feature separation, aiding downstream classifiers.

### Automated Selection of Models

BioFuse automates model selection, identifying the optimal set for each task without manual intervention. It evaluates all 2*^n^* − 1 model combinations, leveraging GPU-accelerated XGBoost to efficiently search this exponential space. Embeddings are fused and rapidly assessed using gradient boosting. GPU acceleration mitigates computational overhead by parallelizing tree construction and handling large feature matrices efficiently. This enables BioFuse to evaluate model combinations in seconds and process larger datasets within minutes, ensuring thorough exploration of the combinatorial space.

An on-disk cache stores previously generated embeddings to avoid redundant computations. BioFuse selects the combination with the highest validation accuracy, ensuring embeddings are tailored to the specific task while maintaining generalisation within the dataset.

## Experimental Setup

### Dataset

As Illustrated in Fig 2, our study uses 12 2D datasets from the MedMNIST+ benchmark [62], a large-scale, MNIST-like collection of standardized biomedical images designed for various machine learning tasks in the medical domain. Key features of the dataset include:

1. **Multi-modal:** MedMNIST+ covers a wide range of biomedical imaging modalities, including X-Ray, OCT, Ultrasound, CT, Histopathology and Electron Microscopy images.
2. **Standardized:** All images are pre-processed into a uniform 224×224 resolution for 2D datasets, with corresponding classification labels. The datasets also come with consistent train-validation-test splits.
3. **Multi-task:** The dataset supports various machine learning tasks, including binary and multi-class classification, ordinal regression, and multi-label classification.
4. **Multi-scale:** MedMNIST includes approximately 708K 2D images, with dataset sizes ranging from 780 to 236,386 samples.

**Fig 2.**
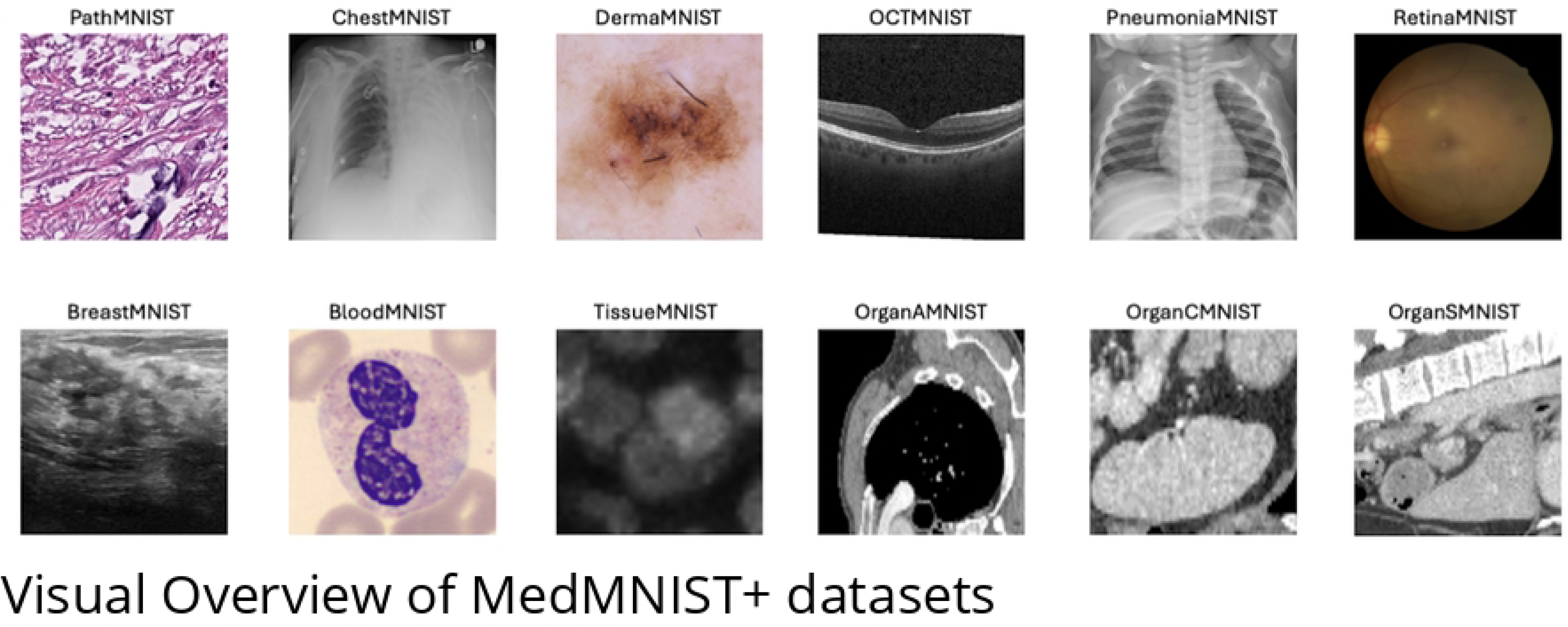
Visual overview of the MedMNIST+ 2D datasets. Sample images at 224×224 resolution showcase the diverse collection spanning multiple medical imaging modalities, illustrating the wide range of diagnostic tasks represented in the benchmark.

The decision to focus on 2D datasets was driven by the use of a simple classifier which is well-suited for 2D image analysis tasks. Table 2 provides an overview of the MedMNIST2D datasets used in this study.

**Table 2.** Overview of MedMNIST+ 2D Datasets.

| Dataset | Data Modality | Tasks(# Classes) | # Samples | # Train/Valid/Test |
| --- | --- | --- | --- | --- |
| PathMNIST [63] | Colon Pathology | Multi-Class (9) | 107,180 | 89,996 / 10,004 / 7,180 |
| ChestMNIST [64] | Chest X-Ray | Multi-Label (14)<br>Binary-Class (2) | 112,120 | 78,468 / 11,219 / 22,433 |
| DermaMNIST [65] | Dermatoscope | Multi-Class (7) | 10,015 | 7,007 / 1,003 / 2,005 |
| OCTMNIST [66] | Retinal OCT | Multi-Class (4) | 109,309 | 97,477 / 10,832 / 1,000 |
| PneumoniaMNIST [66] | Chest X-Ray | Binary-Class (2) | 5,856 | 4,708 / 524 / 624 |
| RetinaMNIST [67] | Fundus Camera | Ordinal Regression (5) | 1,600 | 1,080 / 120 / 400 |
| BreastMNIST [68] | Breast Ultrasound | Binary-Class (2) | 780 | 546 / 78 / 156 |
| BloodMNIST [69] | Blood Cell Microscope | Multi-Class (8) | 17,092 | 11,959 / 1,712 / 3,421 |
| TissueMNIST [70] | Kidney Cortex Microscope | Multi-Class (8) | 236,386 | 165,466 / 23,640 / 47,280 |
| OrganAMNIST [71] | Abdominal CT | Multi-Class (11) | 58,830 | 34,561 / 6,491 / 17,778 |
| OrganCMNIST [71] | Abdominal CT | Multi-Class (11) | 23,583 | 12,975 / 2,392 / 8,216 |
| OrganSMNIST [71] | Abdominal CT | Multi-Class (11) | 25,211 | 13,932 / 2,452 / 8,827 |

Data sourced from [62]

MedMNIST+ enables a comprehensive evaluation of BioFuse across multi-class, binary, and multi-label classification, spanning diverse dataset sizes and class distributions. For consistency, we treat the ordinal regression task (RetinaMNIST) as a multi-class classification problem, a standard approach for handling ordinal data in machine learning [72].

## Experimental Design

### Objectives

Our experiments assess:

1. **Performance Improvement:** Evaluate whether fusing embeddings via vector concatenation outperforms individual models on MedMNIST+ using AUC and Accuracy.
2. **Cross-modal Transfer:** Test if features from one modality (e.g., histopathology) enhance performance when combined with another (e.g., radiographs), measured by AUC and Accuracy.
3. **Generalisation:** Assess BioFuse’s ability to maintain strong performance across diverse biomedical domains using nine foundation models.

These objectives establish BioFuse as a versatile framework for multimodal biomedical imaging, leveraging complementary information to improve classification performance across datasets.

### Hardware Configuration

The experiments were conducted on a server with an NVIDIA A100 GPU (80GB VRAM), an AMD EPYC 7713 CPU (64-core, 128-thread CPU), and 1TB RAM. While the CPU and memory were shared, the GPU was dedicated to BioFuse, ensuring uninterrupted computation for embedding extraction, model fusion, and classification.

## Experimental Procedure

### XGBoost

For internal evaluation (see Fig 1), we used XGBoost [56], a scalable gradient-boosted decision tree algorithm known for its efficiency in binary and multi-class classification. The model was trained on fused embeddings from multiple foundation models.

Although embedding order can influence performance, we found this effect negligible. Optimizing order introduces substantial complexity—2*^n^* − 1 model combinations and *n*! possible orders per combination—making exhaustive search impractical. To maintain computational tractability, we fixed the concatenation order.

Hyperparameters were manually selected following [73], balancing efficiency and accuracy. We set n_estimators = 250, learning_rate = 0.1, and max_depth = 6 to prevent overfitting while capturing biomedical data patterns. The objective function was multi:softprob for multi-class, binary:logistic for binary classification, and a one-versus-rest strategy for multi-label tasks. Training used XGBoost’s GPU acceleration to reduce computation time.

### Training and Evaluation Workflow

Our evaluation of BioFuse follows a structured workflow, as illustrated in Fig 3:

1. **Dataset Preparation:** BioFuse is provided with training and validation sets for each dataset in MedMNIST+ 2D.
2. **Model Selection:** It identifies the optimal combination of foundation models for each dataset (see Fig 1).
3. **Embedding Generation:** Using the selected model combination, BioFuse generates training and validation embeddings and returns a configured BioFuseModel (embedding generator).
4. **Classifier Selection and Tuning:** For the final evaluation, we use XGBoost, tuning its hyperparameters on training embeddings and selecting the configuration that achieves the highest validation accuracy.
5. **Test Embedding Generation:** The trained BioFuseModel generates embeddings for the test set, ensuring consistency with the training and validation process.
6. **Final Evaluation:** The tuned XGBoost model (from Step 4) is evaluated on the test set embeddings to assess overall performance.

**Fig 3.**
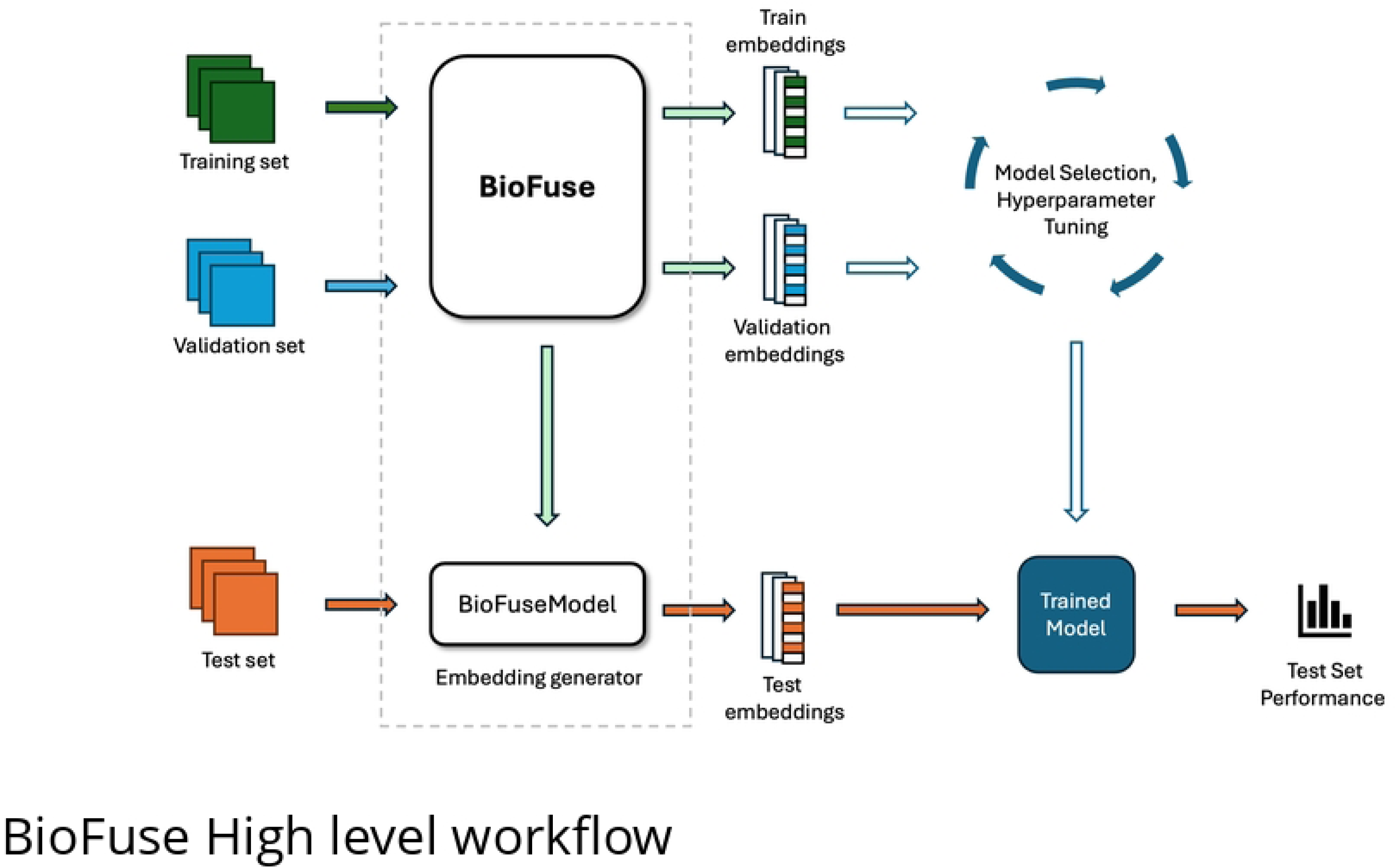
High-level workflow of BioFuse within a typical machine learning pipeline. BioFuse serves as a sophisticated embedding generator, accepting training and validation sets as inputs. As outputs, BioFuse provides embeddings for both the training and validation sets, along with a configured BioFuseModel. This BioFuseModel acts as an embedding generator for subsequent use on the test set.

To ensure a comprehensive final evaluation on the test sets, we conducted independent hyperparameter tuning for each dataset in the MedMNIST+ benchmark (as described in Step 4). Details on the specific hyperparameters are provided in Appendix B (Table 6)for reproducibility.

Performance was measured using test AUC and accuracy, with results averaged over three independent runs to enhance robustness.

### Evaluation Metrics

We employ two complementary metrics in our evaluation framework:

**Accuracy** is used for internal model selection during BioFuse’s automated search process:

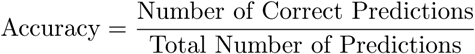

**Area Under the ROC Curve (AUC-ROC)** serves as our primary reporting metric for final test set performance. This choice aligns with MedMNIST+ benchmarks and provides a threshold-independent assessment of model quality. AUC-ROC represents the probability that a randomly chosen positive example ranks higher than a negative one [74], with values ranging from 0.5 (random performance) to 1.0 (perfect classification).

For comprehensive comparison, we report both AUC-ROC and accuracy in our final results tables.

## Results

### Overview

On the MedMNIST+ benchmark, BioFuse outperforms several existing methods, achieving the highest test AUC on 5 of 12 datasets and near-SOTA performance on six more. It also achieves the best test accuracy on 2 of 12 datasets and near-SOTA accuracy on another 4 (see Table 3). These results highlight the effectiveness of embedding fusion from diverse foundation models in improving performance across a broad range of biomedical image classification tasks.

**Table 3.**
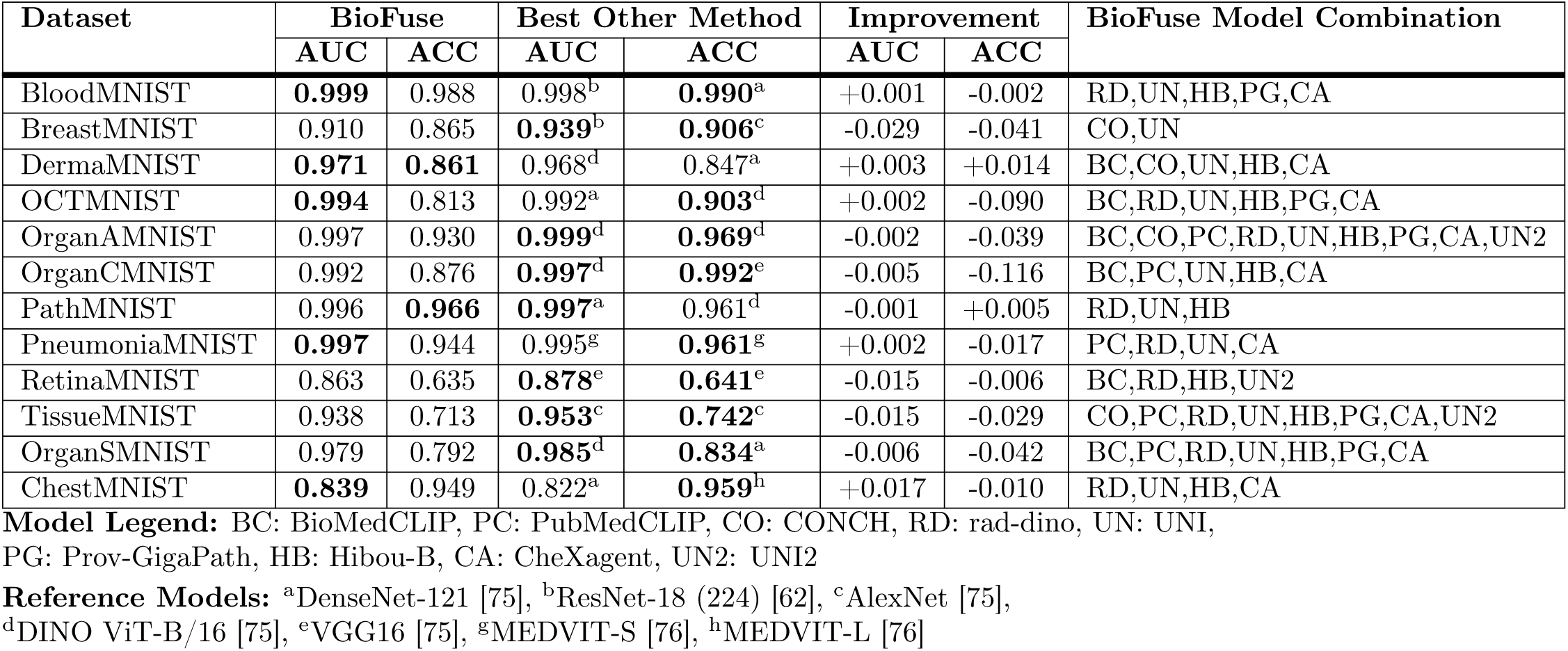
Performance Comparison Between BioFuse and Best Existing Methods on MedMNIST+. Bold values indicate the best performance for each dataset.

A particularly interesting finding is the versatility of histopathology and radiology foundation models. Histopathology models UNI, Hibou-B, CONCH and Prov-GigaPath and radiology models CheXagent and rad-dino consistently appear in the top-performing model combinations, even when applied to modalities different from their pre-training data. This cross-modality effectiveness suggests these models learn generalisable features useful across various biomedical imaging tasks.

## Cross-Modal Transfer Performance

### Transfer to other medical imaging modalities

We observed notable cross-modal transfer capabilities across both histopathology and radiology models when evaluating each model independently across the datasets, as illustrated in Fig 4 and Fig 5. This single-model evaluation used fixed hyperparameters for XGBoost as described in Experimental Procedure. As a baseline, we included CLIP [32], a general-purpose vision-language foundation model, to compare performance against biomedical foundation models. CLIP’s performance was generally low across all datasets, with the notable exception of OrganAMNIST, where it ranked in the top 3 for both accuracy and AUC.

**Fig 4.**
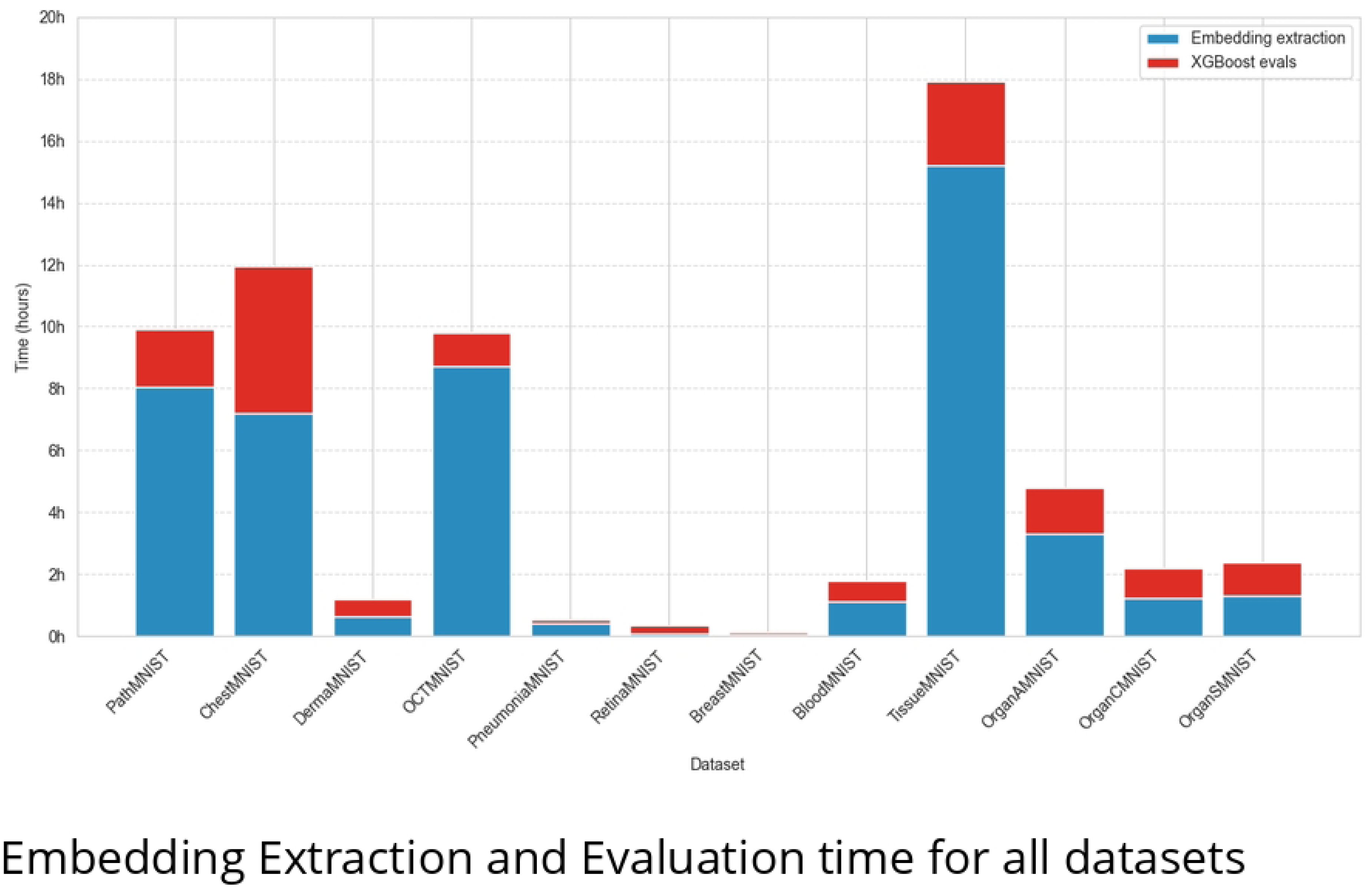
Heatmap of test AUC, showing single model performance across MedMNIST+ datasets. CLIP is included as a baseline general-purpose foundation model for comparison. White box indicates the top-performer while red boxes highlight the 2nd and 3rd top performers for each dataset; More than three models may be highlighted if they share identical values.

**Fig 5.**
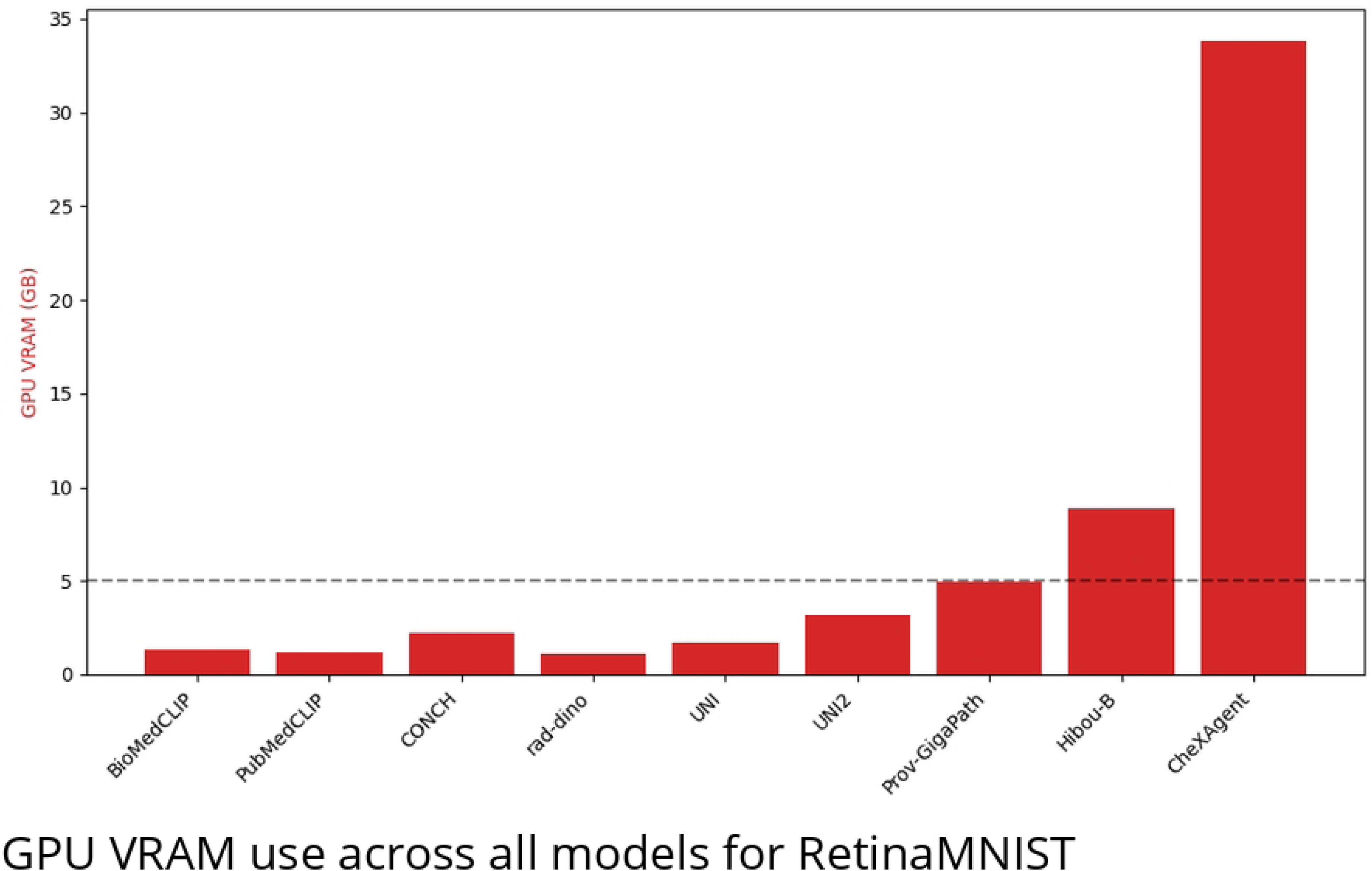
Heatmap showing test accuracy, showing single model performance across MedMNIST+ datasets and ImageNet-1K (top-1). CLIP is included as a baseline general-purpose foundation model for comparison. White boxes indicate top performers, while red boxes highlight second and third best performers for each dataset. When multiple models achieve identical performance, more than three boxes may be highlighted.

Fig 4 shows that Prov-GigaPath, Hibou-B, and UNI demonstrated strong performance in various radiology datasets. Prov-GigaPath achieved the highest AUC on BreastMNIST and ranked among the top-3 for OrganAMNIST. Hibou-B secured top-3 AUC in BreastMNIST, OrganCMNIST and OrganSMNIST. UNI excelled in CT and X-ray tasks, achieving the highest AUC for OrganSMNIST and top-3 in PneumoniaMNIST. CheXagent, primarily trained on chest X-rays, demonstrated impressive cross-modal capabilities, achieving top performance in DermaMNIST and ranking among the top-3 for BloodMNIST, TissueMNIST, and RetinaMNIST.

Fig 5 corroborates this trend, with Hibou-B achieving top-3 accuracy on both OrganCMNIST and OrganSMNIST. Prov-GigaPath demonstrated excellence in radiology tasks, achieving top accuracy for BreastMNIST and ranking in top-3 for OrganAMNIST, OrganCMNIST, OrganSMNIST, and PneumoniaMNIST. UNI showed remarkable performance, tying for top accuracy in BreastMNIST and leading in OrganSMNIST. CheXagent’s versatility is again evident, achieving top accuracy in both DermaMNIST and RetinaMNIST, while securing top-3 positions in BloodMNIST and TissueMNIST.

### Transfer to Natural Images

To comprehensively evaluate the generalisation capabilities of medical foundation models, we assess their performance on the ImageNet-1K [77] test set, a benchmark dataset widely used in computer vision. While we don’t expect medical models to match the performance of models trained specifically on natural images, their performance provides valuable insights into their general visual understanding capabilities. Surprisingly, some medical models demonstrate remarkable transfer ability, with Prov-GigaPath and CheXAgent achieving 78.2% and 66.0% Top-1 accuracy respectively, outperforming CLIP (58.1%) despite CLIP being trained on 400M natural image-text pairs. Table 4 presents the complete ImageNet-1K test set performance results.

**Table 4.** Performance Comparison of Biomedical Foundation Models on ImageNet-1K. Bold values indicate best performance.

| Model | Top-1 ACC | Top-5 ACC | Pre-training Dataset Size |
| --- | --- | --- | --- |
| CoCa | **0.910** | - | 4.8B (3B labeled images + 1.8B image-text pairs) |
| Prov-GigaPath | 0.782 | 0.944 | 1.3B tiles (171K WSIs) |
| CheXAgent | 0.660 | 0.908 | 6.1M I-Q-A triplets (1.1M CXRs) |
| CLIP | 0.581 | 0.870 | 400M image-text pairs |
| PubMedCLIP | 0.466 | 0.772 | 81K image-text pairs |
| UNI2 | 0.274 | 0.517 | 100M tiles (100K WSIs) |
| BioMedCLIP | 0.240 | 0.502 | 15M image-text pairs |
| UNI | 0.240 | 0.479 | 100M tiles (100K WSIs) |
| Hibou-B | 0.220 | 0.444 | 512M tiles (1M WSIs) |
| CONCH | 0.158 | 0.347 | 1.17M image-caption pairs |
| rad-dino | 0.114 | 0.268 | 838K images |

## Discussion

### Key Findings

Based on the comprehensive results in Table 3, we would like to highlight some important observations:

- **State-of-the-Art Performance:** BioFuse demonstrates exceptional performance on the MedMNIST+ benchmark, achieving new stateof-the-art (SOTA) scores across multiple datasets. Specifically, it achieves the highest AUC in five datasets and the best accuracy in two datasets. These improvements, achieved in a highly competitive benchmark, underscore BioFuse’s ability to extract and combine more informative features from multiple pre-trained models.
- **Strong Performance on DermaMNIST:** A notable improvement was observed on the DermaMNIST dataset, where BioFuse achieved state-of-the-art performance in both AUC and accuracy. Despite the SOTA AUC already being at 0.97, BioFuse managed to deliver a further performance improvement in AUC and 1.4% in Accuracy. This demonstrates BioFuse’s ability to improve performance even on tasks where the margin for improvement is small.
- **Near SOTA AUC performance:** BioFuse demonstrated highly competitive performance across multiple datasets, achieving AUC scores within 1.5% of state-of-the-art methods. This consistent performance was observed across various CT imaging tasks (OrganAMNIST, OrganCMNIST, OrganSMNIST) and microscopy datasets (TissueMNIST, PathMNIST). While BioFuse did not achieve the highest test accuracy for most datasets, it notably achieved SOTA performance on PathMNIST. The minimal gap in AUC scores compared to best-performing methods suggests BioFuse maintains robust class separation capabilities across these diverse medical imaging tasks.
- **AUC Superiority in Lower Accuracy Datasets:** For datasets such as OCTMNIST, PneumoniaMNIST, and ChestMNIST, BioFuse’s test accuracy lagged behind existing models, but it outperformed them in AUC scores. Specifically, BioFuse achieved an AUC of 0.994 (vs. 0.992) for OCTMNIST, 0.997 (vs. 0.995) for PneumoniaMNIST, and 0.839 (vs. 0.822) for ChestMNIST, showing that even when accuracy lags, BioFuse’s embeddings allow the model to make confident and accurate class distinctions.
- **Efficacy of Model Ensembles:** Across all datasets, the best-performing combinations involved multiple foundation models. No

single model was able to outperform the ensembles, underscoring the importance of model diversity in feature representation. This finding suggests that the combined outputs from different models allow BioFuse to better capture complementary information across multiple modalities, resulting in more robust performance across a variety of biomedical imaging tasks.

### Cross-Modal Transfer

Biomedical foundation models exhibited remarkable cross-modal transfer capabilities, often excelling in tasks well beyond their original training domains. This generalisation likely stems from shared visual feature representations that transcend specific imaging modalities. Notably, models trained on a single modality (such as histopathology-specific models) demonstrated stronger cross-modal transfer than those trained on multiple modalities, suggesting a potential benefit to focused, modality-specific pre-training for developing generalisable representations.

Histopathology models — particularly UNI, Hibou-B, and Prov-GigaPath — demonstrated exceptional cross-modal performance, likely due to their Vision Transformer-based architectures. These models develop capacity to process multi-scale visual features during pre-training, from micron-scale cellular details to millimeter-scale tissue structures. This multi-scale capability appears particularly transferable across modalities, as the feature extraction mechanisms that identify cellular boundaries and tissue organization in histopathology may transfer effectively to detecting analogous structural patterns in retinal images and radiological scans, despite differences in visual appearance.

CheXagent’s sophisticated architecture, which integrates an 8-billion parameter clinical language model with a vision encoder and vision-language bridge, may contribute to its strong cross-modal adaptability. Pre-trained on 6 million instruction-image-answer triplets across 65 diverse datasets, this model appears to develop both modality-specific and modality-agnostic feature representations. The language component potentially functions as a semantic intermediary, facilitating knowledge transfer between distinct imaging domains. This architectural advantage correlates with CheXagent’s impressive performance in non-radiological tasks such as dermatology (AUC 0.960) and microscopy (AUC 0.999).

While our findings demonstrate substantial cross-modal capabilities, this transfer is not universal across all biomedical tasks. We observed performance decreases in highly specialized domains such as CT scan interpretation, where OrganAMNIST and OrganCMNIST showed the widest performance gaps between specialized and cross-modal models. These limitations suggest that effective cross-modal transfer depends on both model architecture and the inherent similarity between source and target domains. Future work should explore these boundaries systematically, potentially guiding the development of more universally transferable biomedical foundation models.

### Computational Considerations

Understanding BioFuse’s computational demands is essential for practical implementation. Following [78], we analyzed total response time as a realistic computational performance metric. Evaluating BioFuse across 12 datasets required **47h 19m** for embedding extraction and **15h 35m** for model evaluation, totaling **62h 54m**.

Fig 6 presents a detailed breakdown of computation time across all MedMNIST+ datasets. The distribution reveals significant performance variation based on dataset characteristics. TissueMNIST demanded the most processing time (17.8 hours), primarily due to its large size. In contrast, smaller datasets like RetinaMNIST and BreastMNIST required less than 0.5 hours each for complete processing.

**Fig 6.**
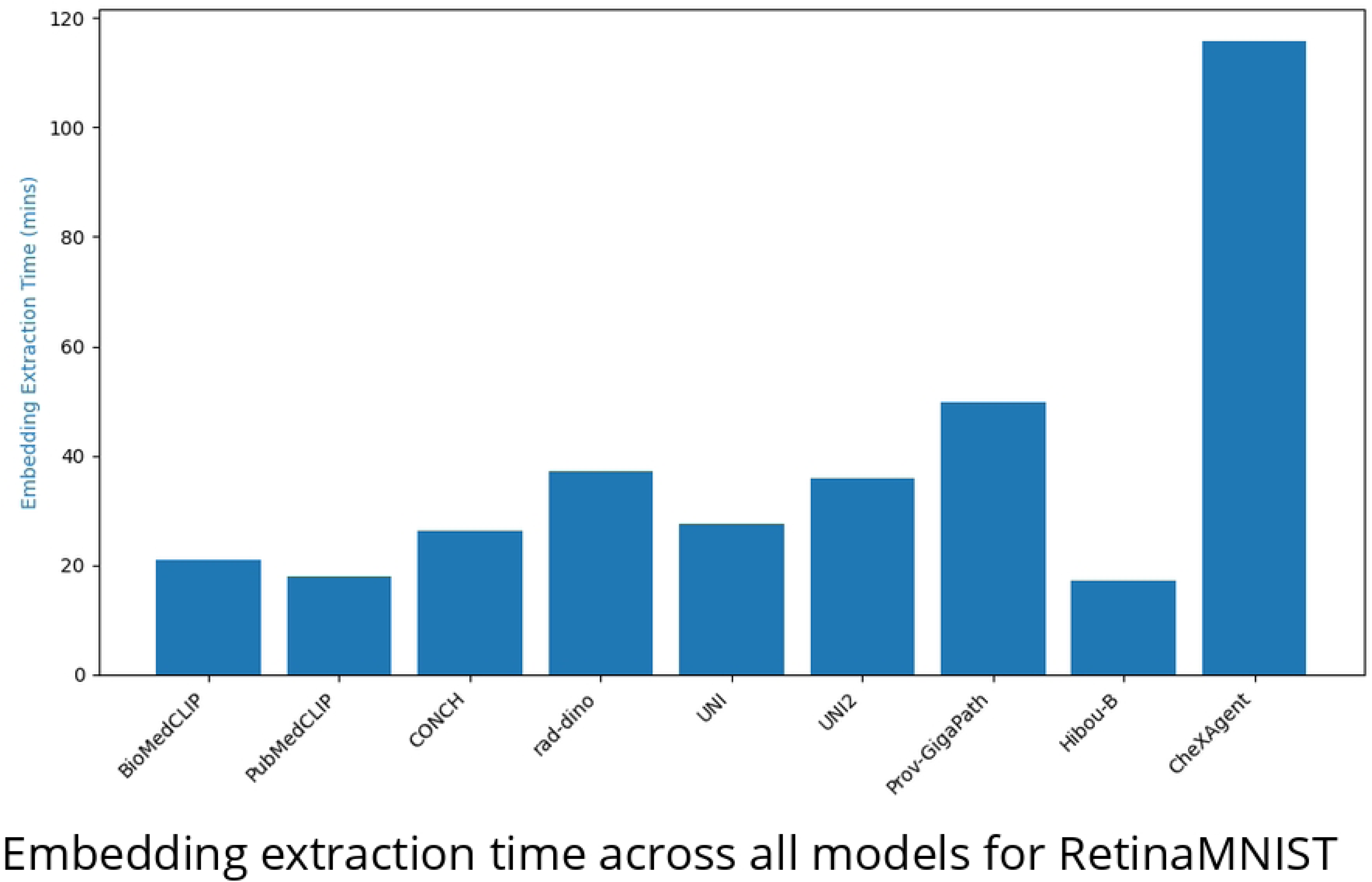
Stacked bar chart showing the total evaluation time for each dataset. The chart is divided into embedding extraction and XGBoost evaluation phases. The blue section represents the time spent on embedding extraction, while the red section indicates the time for model training and validation using XGBoost.

The data reveals that embedding extraction represents the dominant computational bottleneck, accounting for approximately 75% of total processing time. This bottleneck scales primarily with the size of the dataset, as clearly demonstrated by TissueMNIST (236,386 samples) which require the longest extraction time, followed by OCTMNIST (109,309 samples) and PathMNIST (107,180 samples). The relationship between computation time and dataset size is largely linear for each model, as all images are standardized to 224×224 resolution.

In particular, ChestMNIST exhibits an unusually high proportion of time devoted to XGBoost evaluation relative to other datasets (approximately 40% of total processing time versus typical 20-25%). This anomaly stems primarily from ChestMNIST’s multi-label classification nature with 14 possible labels, substantially increasing the complexity of the classification task compared to binary or single-label multi-class problems.

Despite using high-dimensional concatenated embeddings, XGBoost training and evaluation remained computationally efficient in most cases, typically requiring less than 25% of the total computation time. This reinforces XGBoost’s reputation for handling high-dimensional data effectively. Our analysis of GPU memory requirements showed that 7 of 9 foundation models operated within 5GB of VRAM, making them suitable for consumer-grade GPUs (Appendix C Fig 7). However, CheXagent, with 8 billion parameters, required 33.8GB VRAM, necessitating server-grade GPU hardware (Appendix C Fig 8). Although memory requirements scale with data set size, the relative differences between models remain consistent. These resource considerations are particularly important for researchers planning to implement BioFuse in environments with limited computational infrastructure.

**Fig 7.**
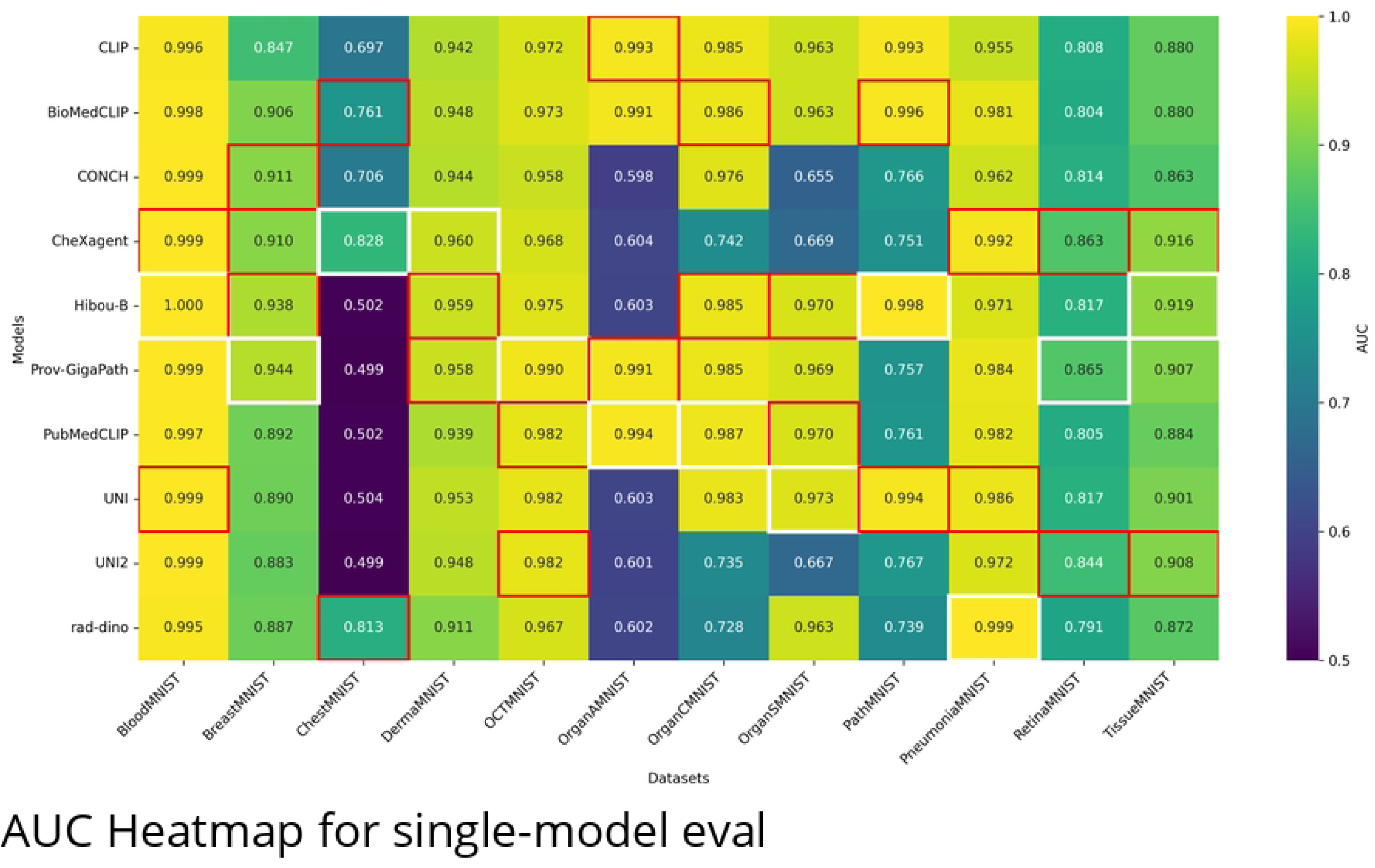
GPU VRAM usage (in GB) for each model when processing the RetinaMNIST dataset. CheXagent has the highest memory demand, requiring more than 33 GB of VRAM for the 1,080 training samples.

**Fig 8.**
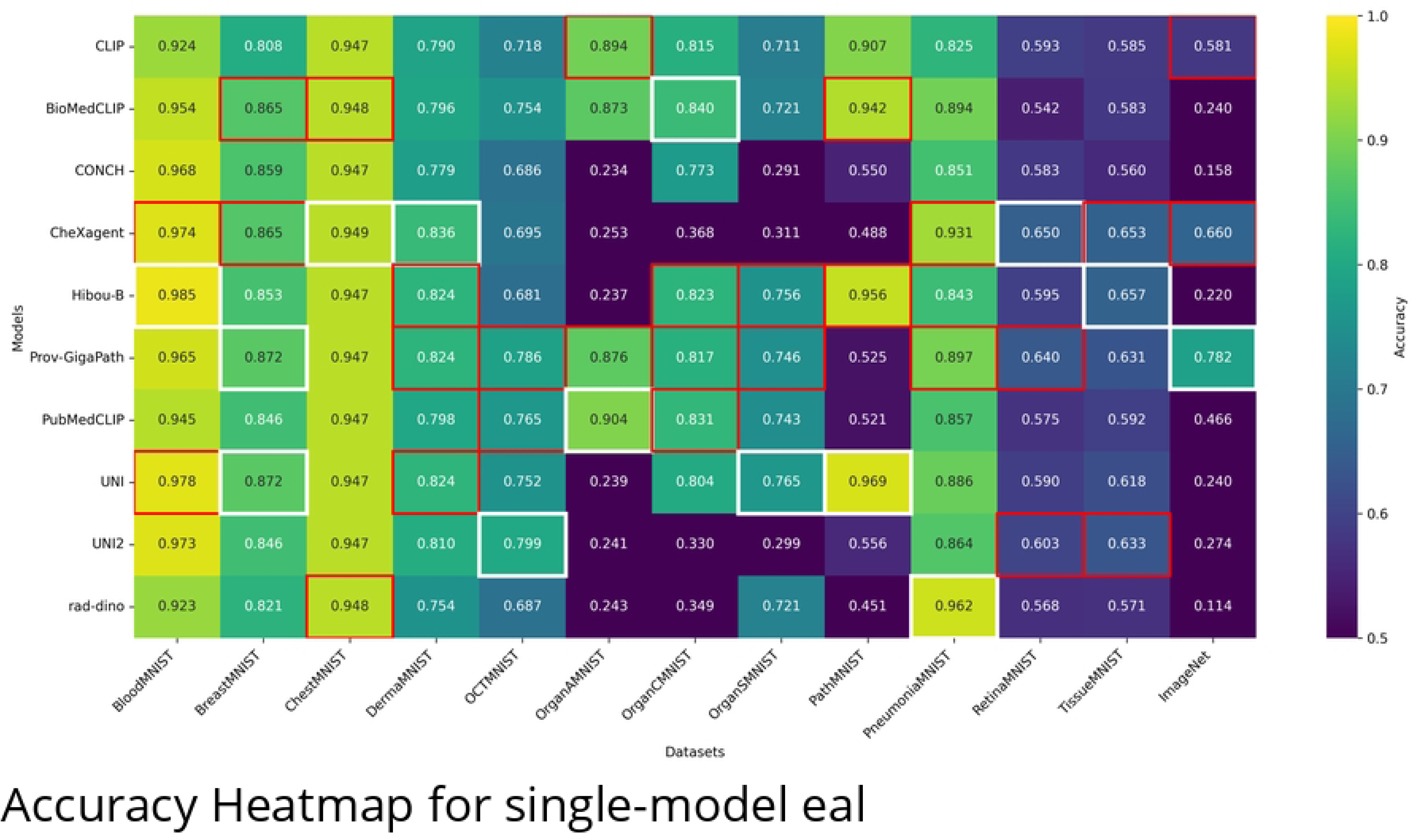
Embedding extraction time (in mins) for each model when processing the RetinaMNIST dataset. CheXagent takes the longest extraction time, significantly exceeding that of other models for the 1,080 training samples.

To support reproducibility and eliminate the need for time-intensive embedding extraction, we provide the complete embedding cache via Zenodo for MedMNIST+ [79] and ImageNet-1K [80].

### Limitations and Future Directions

While BioFuse demonstrates strong potential, several limitations must be addressed in future research:

**Evaluation Scope and Task Generalisation:** BioFuse is currently evaluated only on classification tasks (binary, multi-class, and multi-label) using MedMNIST+. Testing on broader task types (e.g., segmentation, object detection, regression) and real-world clinical datasets would provide more comprehensive validation. Future work should explore BioFuse’s applicability to time-series medical data and multimodal clinical records, potentially expanding its utility in diagnostic and prognostic applications. **Dependency on Pre-trained Models:** BioFuse’s performance inherently depends on the quality and diversity of its foundation models. If these models lack generalisability or introduce biases, these issues may propagate through the fusion process. Expanding the model pool to include diverse biomedical domains (e.g., genomics, MRI, ultrasound, and clinical text) could mitigate this limitation and enable truly multimodal integration beyond imaging.

**Computational Overhead:** Extracting embeddings from multiple models and performing exhaustive grid searches requires significant GPU resources, particularly when using advanced models like CheXagent (33.8GB VRAM). Developing more efficient search strategies that can identify optimal model combinations without exhaustively evaluating all possibilities represents another promising direction.

**Vector Concatenation Limitations:** The current approach can lead to high-dimensional embeddings with redundancy, potentially increasing overfitting risks on smaller datasets. Exploring alternative fusion strategies, such as attention-based fusion, knowledge distillation, or learned projection spaces, may improve efficiency and reduce dimensionality issues while preserving information content.

**Interpretability Challenges:** Combining multiple models reduces transparency in how predictions are formed, which is critical for clinical applications. Integrating explainability techniques like saliency maps, feature attribution, or attention visualization could improve interpretability, helping clinicians understand feature contributions from different models. This remains essential for responsible deployment in healthcare settings.

By addressing these areas through interdisciplinary collaboration between AI researchers, clinicians, and domain experts, BioFuse can evolve into a more efficient, interpretable, and generalisable tool for multimodal biomedical research and clinical applications.

## Conclusion

The integration of diverse foundation models represents a promising frontier for advancing biomedical imaging analysis. In this work, we introduced BioFuse, a novel framework that systematically fuses embeddings from multiple biomedical foundation models to generate optimised representations for downstream tasks. Evaluated across 12 diverse imaging modalities in the MedMNIST+ benchmark, BioFuse with XGBoost classification outperformed existing methods, achieving the highest AUC in five datasets and maintaining near-SOTA performance in most others. Notably, it demonstrated exceptional performance in dermatology classification (DermaMNIST) and revealed unexpected cross-modal transfer capabilities in histopathology and radiology models like UNI, Hibou-B, Prov-GigaPath, and CheXagent.

These results highlight the benefits of leveraging multiple pre-trained models rather than relying on a single foundation model. BioFuse’s ability to automatically identify and integrate complementary representations from diverse models suggests significant potential for healthcare applications requiring comprehensive image interpretation across modalities. The framework’s extensible architecture ensures adaptability to future foundation models as they emerge.

While demonstrating clear advantages, BioFuse faces challenges, including computational overhead from grid search and potential redundancy in concatenated embeddings. Future work should explore more efficient fusion strategies, expand applications beyond classification to segmentation and detection tasks, and incorporate interpretability mechanisms essential for clinical adoption and regulatory approval.

By harnessing the collective strengths of multiple foundation models through a systematic approach to embedding fusion, BioFuse not only improves performance on benchmark tasks but also opens new avenues for cross-modal knowledge transfer in biomedical imaging. This contribution moves us closer to more reliable and comprehensive AI-assisted medical decision-making systems that can integrate information across the diverse imaging modalities encountered in clinical practice.

## Funding

This research was supported by the School of Computer Science at the University of St Andrews through a PhD studentship. The funder had no role in study design, data collection and analysis, decision to publish, or preparation of the manuscript.

## Acknowledgments

We thank the School of Computer Science at the University of St Andrews for providing computational resources and support. We also appreciate the IT team for their technical assistance.

## Competing Interests

The authors have declared that no competing interests exist.

## Ethics Statement

This research was conducted under ethical approval from the institutional review board (approval code CS17485).

## Footnotes

1 Code: https://github.com/mnhcorp/biofuse

## Supporting information

### S1 Appendix. BioFuse API Specification

#### Typical workflow

BioFuse’s workflow is split into 5 stages:

1. Initialize BioFuse with selected foundation models.
2. Generate BioFuse embeddings for training and validation datasets.
3. Train and tune downstream model using the BioFuse embeddings.
4. Generate test embeddings using the BioFuseModel embedding generator
4. Evaluate the performance of the final model using test embeddings.

#### High-level API

##### 1. Initialize BioFuse

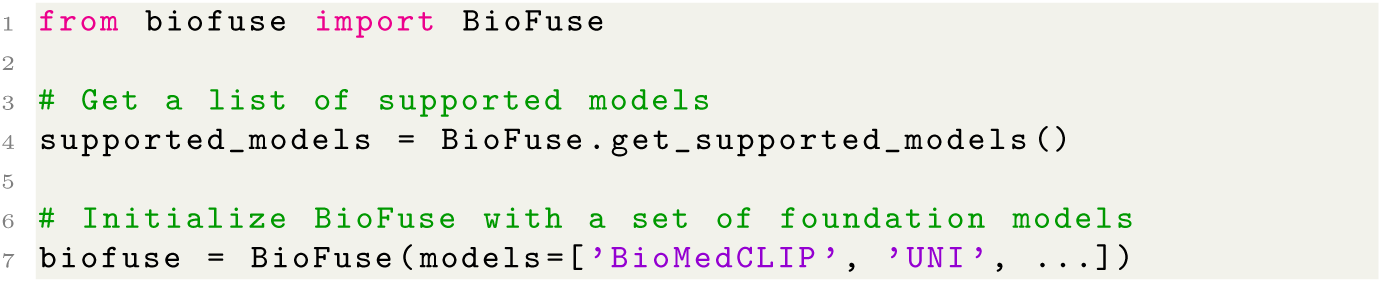

##### 2. Generate BioFuse Embeddings

Generate embeddings from the training and validation sets (X,y) pairs along with a BioFuseModel that can be used for generating embeddings for new data. BioFuse also expects the task type, currently supporting binary, multi-class and multi-label classification.

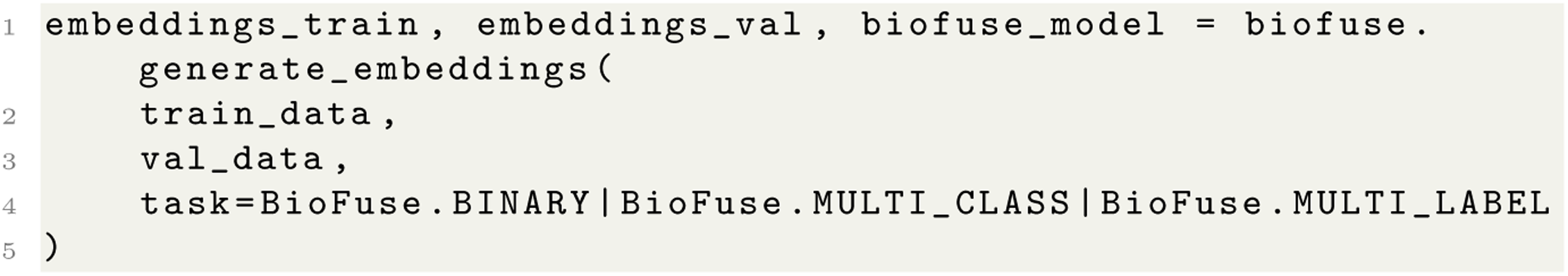

##### 3. Generate BioFuse Embeddings for new data using BioFuseModel

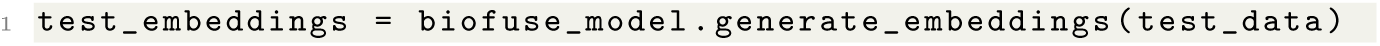

### Example Usage

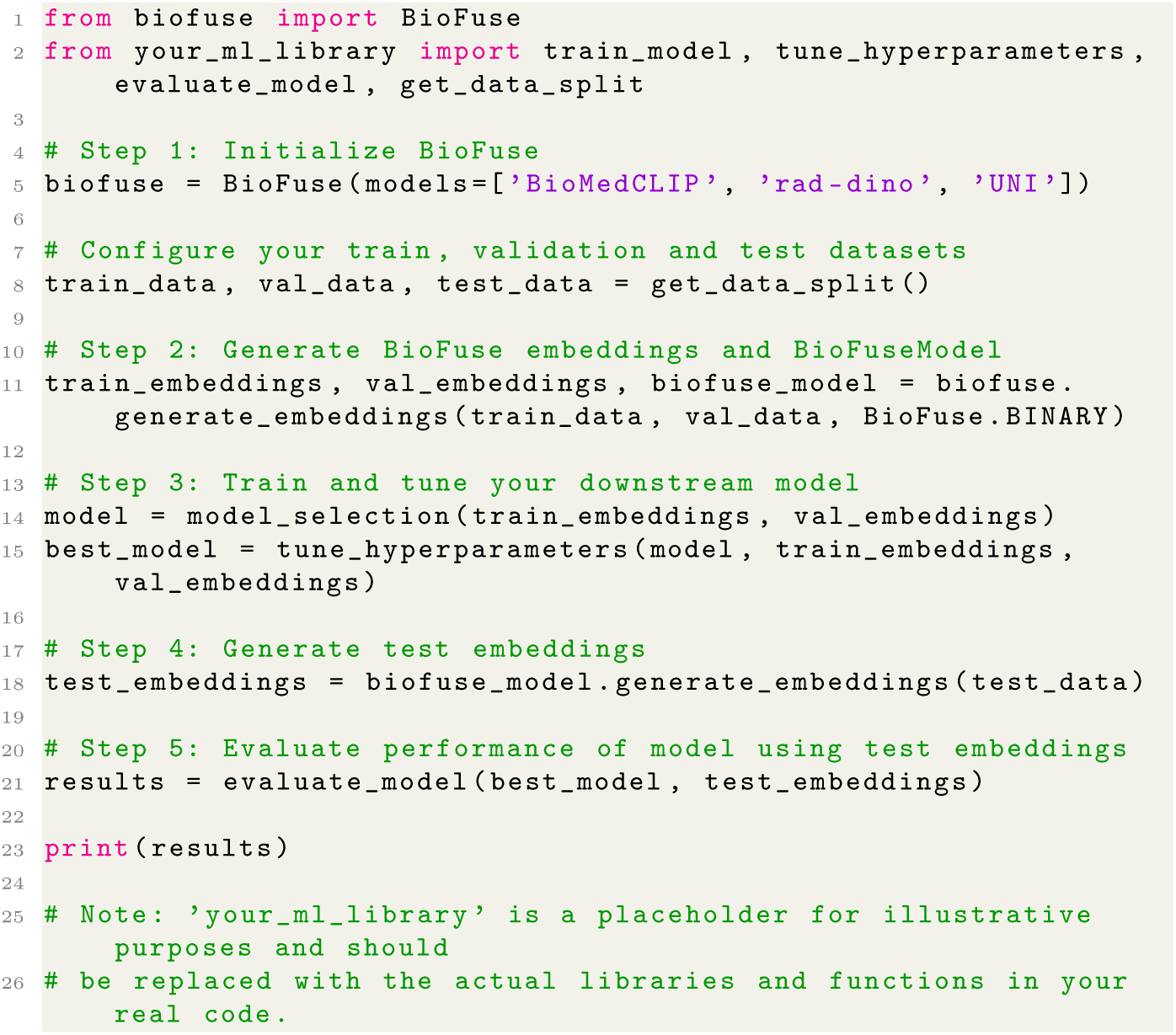

### S2 Appendix. Hyperparameter Tuning Configurations for XGBoost

#### Overview

This appendix provides the specific hyperparameter configurations used for XGBoost on each dataset in MedMNIST+. Given XGBoost’s extensive hyperparameter tuning space, we employed the Bayesian optimization method through the Weights Biases (wandb) platform. This approach allowed us to efficiently navigate the vast hyperparameter landscape and identify optimal configurations for each dataset, balancing the exploration of the parameter space with the exploitation of promising regions.

#### Hyperparameter Tuning setup

Table 5 presents the hyperparameters considered and their respective search ranges. For each parameter, we selected ranges that encompass both conservative and aggressive values to thoroughly explore the model’s behavior.

**Table 5.** XGBoost Hyperparameters and Their Search Ranges.

| Parameter | Range | Description |
| --- | --- | --- |
| learning_rate | [0.01, 0.03, 0.05, 0.1, 0.2, 0.3] | Step size shrinkage used to prevent overfitting. Lower values make the model more robust but slower to compute |
| n_estimators | [100, 250, 500, 750, 1000, 1500, 1750, 2000, 2250, 2500] | Number of boosting rounds or trees to be built |
| max_depth | [3, 6, 9, 12, 15] | Maximum depth of each tree. Larger values increase model complexity |
| max_leaves | [0, 31, 63, 127, 255] | Maximum number of leaves in each tree. 0 indicates no limit |
| min_child_weight | [1, 3, 5, 7] | Minimum sum of instance weight needed in a child node |
| subsample | [0.6, 0.8, 1.0] | Fraction of samples used for training each tree |
| colsample_bytree | [0.6, 0.8, 1.0] | Fraction of features used for training each tree |
| reg_alpha | [0, 0.1, 1, 10] | L1 regularization term on weights to prevent overfitting |
| reg_lambda | [0, 0.1, 1, 10] | L2 regularization term on weights to prevent overfitting |
| max_bin | [128, 256, 512] | Maximum number of discrete bins for continuous features |

**Table 6.** Optimal XGBoost Hyperparameter Configurations for MedMNIST+ 2D.

| Dataset | max depth | colsample bytree | learning rate | max bin | max leaves | child weight | num estimators | reg alpha | reg lambda | sub sample |
| --- | --- | --- | --- | --- | --- | --- | --- | --- | --- | --- |
| OrganAMNIST | 9 | 1.0 | 0.2 | 128 | 31 | 7 | 1500 | 10.0 | 1.0 | 1.0 |
| OrganSMNIST | 9 | 1.0 | 0.1 | 256 | 255 | 1 | 1500 | 1.0 | 0.1 | 1.0 |
| OrganCMNIST | 3 | 0.8 | 0.1 | 512 | 31 | 5 | 1500 | 1.0 | 1.0 | 0.6 |
| PathMNIST | 15 | 0.8 | 0.1 | 128 | 31 | 1 | 1500 | 1.0 | 10.0 | 0.6 |
| BloodMNIST | 15 | 0.6 | 0.3 | 128 | 255 | 3 | 1500 | 1.0 | 10.0 | 0.8 |
| OCTMNIST | 15 | 0.8 | 0.2 | 512 | 255 | 3 | 1500 | 0.0 | 0.1 | 1.0 |
| BreastMNIST | 3 | 1.0 | 0.01 | 512 | 127 | 1 | 500 | 10.0 | 10.0 | 1.0 |
| ChestMNIST | 15 | 1.0 | 0.03 | 128 | 127 | 7 | 1000 | 10.0 | 10.0 | 1.0 |
| TissueMNIST | 12 | 1.0 | 0.3 | 512 | 63 | 5 | 1500 | 0.1 | 0.1 | 1.0 |
| DermaMNIST | 15 | 0.6 | 0.05 | 512 | 255 | 3 | 750 | 10.0 | 10.0 | 1.0 |
| PneumoniaMNIST | 12 | 1.0 | 0.1 | 128 | 255 | 7 | 1000 | 10.0 | 0.0 | 1.0 |
| RetinaMNIST | 12 | 1.0 | 0.1 | 512 | 127 | 5 | 500 | 1.0 | 1.0 | 1.0 |

#### Dataset specific configurations

After conducting an extensive hyperparameter search, we identified the optimal configurations for each dataset in the MedMNIST collection. Table 6 presents these optimised hyperparameter settings, which show interesting variations across different medical imaging tasks.

### S3 Appendix. Computation Time

#### S4 Appendix. Pretraining Datasets for Biomedical Foundation Models

The table below details the datasets used in the pretraining of biomedical foundation models used by BioFuse. This ensures transparency regarding the sources and content of each dataset, clarifying any overlaps or distinctions between the training data of individual models. Such information is critical for evaluating dataset independence, particularly in relation to BioFuse’s target datasets, like MedMNIST.

**Table 7.** Pretraining Datasets for Models in BioFuse.

| Model | Dataset(s) Used | Dataset Details |
| --- | --- | --- |
| UNI [39] | Mass-100K | Over 100M tissue patches from 100,426 WSIs covering 20 tissue types, sourced from MGH, BWH, and GTEx. |
| CONCH [47] | Histopathology Image-Caption Pairs | 1.17M pairs, combining educational resources and PubMed Central, limited to human histopathology. |
| Hibou-B [42] | Proprietary WSI Dataset | 1M WSIs from various tissue types, including 936,441 H&E and 202,464 non-H&E slides, and veterinary cases. |
| Prov-GigaPath [41] | Prov-Path | 1.3B 256×256 tiles from 171,189 pathology slides across 31 tissue types from 30,000+ patients. |
| CheXagent [46] | CheXinstruct | 6M image-instruction-answer triplets from 65 CXR datasets, including MIMIC-CXR, PadChest, and BIMCV-COVID-19. |
| RAD-DINO [43] | Multi-CXR | 838,000 CXR images from BRAX, MIMIC-CXR, NIH-CXR, PadChest, and private outpatient studies. |
| BioMedCLIP [44] | PMC-15M | 15M image-text pairs from 4.4M PubMed Central articles, covering 30 biomedical image types. |
| PubMedCLIP [45] | ROCO | 80,000 image-text pairs from PubMed, including ultrasound, X-ray, MRI, and other modalities across body regions. |

MGH - Massachusetts General Hospital, BWH - Brigham and Women’s Hospital, GTEx - Genotype-Tissue Expression Consortium

#### S5 Appendix. MedMNIST+ Dataset Details

The MedMNIST+ collection provides a diverse set of labeled medical imaging datasets across various diagnostic domains, designed to support machine learning and deep learning research in healthcare. The table below outlines the source datasets, imaging modalities, and primary diagnostic categories for each MedMNIST+ subset, highlighting the origins and clinical relevance of each dataset. Importantly, this table serves as evidence that none of the datasets in MedMNIST+ overlap with the pretraining datasets used in the biomedical foundation models integrated within BioFuse.

**Table 8.** MedMNIST Datasets and Their Sources.

| Dataset | Source Dataset(s) | Description |
| --- | --- | --- |
| PathMNIST [63] | Colorectal Cancer Histology Dataset | H&E-stained histology images from 86 slides, comprising over 100,000 patches, representing tissue types associated with colorectal cancer. |
| OCTMNIST [66] | Retinal OCT Dataset | OCT images for retinal disease diagnosis, including macular degeneration and diabetic macular edema, from institutions like Shiley Eye Institute. |
| PneumoniaMNIST [66] | Guangzhou Women and Children’s Medical Center | 5,232 pediatric chest X-ray images labeled to diagnose bacterial and viral pneumonia. |
| ChestMNIST [64] | ChestX-ray8 | Contains 108,948 frontal-view X-ray images of thoracic diseases such as pneumonia, cardiomegaly, and pneumothorax, derived from NIH. |
| DermaMNIST [65] | HAM10000 | 10,015 dermatoscopic images from Queensland and Vienna clinics, covering seven skin lesion categories including melanoma and nevi. |
| BloodMNIST [69] | Hospital Clinic of Barcelona Blood Cells | 17,092 peripheral blood cell images, categorized into eight cell types, captured using the CellaVision DM96 analyzer. |
| BreastMNIST [68] | Breast Ultrasound Dataset | 780 breast ultrasound images from Baheya Hospital in Cairo, categorized as normal, benign, or malignant. |
| RetinaMNIST [67] | DeepDRiD | Fundus images from 500 patients, used for grading diabetic retinopathy and image quality estimation. |
| TissueMNIST [70] | Broad Bioimage Benchmark Collection (BBBC) | High-throughput microscopy images of cells and organisms for image-analysis benchmarking. |
| Organ{A,S,C}MNIST [71] | Liver Tumor Segmentation Benchmark (LiTS) | CT images focused on liver tumors, collected from multiple institutions, labeled for primary and secondary tumor segmentation. |

## Notes

### Competing Interest Statement

The authors have declared no competing interest.

